# Adult neurogenesis mediates forgetting of multiple types of memory in the rat

**DOI:** 10.1101/2021.03.22.436536

**Authors:** Gavin A. Scott, Dylan J. Terstege, Andrew J. Roebuck, Kelsea A. Gorzo, Alex P. Vu, John G. Howland, Jonathan R. Epp

## Abstract

The formation and retention of hippocampus-dependent memories is impacted by neurogenesis, a process that involves the production of new neurons in the dentate gyrus of the hippocampus. Recent studies demonstrate that increasing neurogenesis after memory formation induces forgetting of previously acquired memories. Neurogenesis-induced forgetting was originally demonstrated in mice, but a recent report suggests that the same effect may be absent in rats. Although a general species difference is possible, other potential explanations for these incongruent findings are that memories which are more strongly reinforced become resilient to forgetting or that perhaps only certain types of memories are affected. Here, we investigated whether neurogenesis-induced forgetting occurs in rats using several hippocampal dependent tasks including contextual fear conditioning (CFC), the Morris Water Task (MWT), and touchscreen paired associates learning (PAL). Neurogenesis was increased following training using voluntary exercise for 4 weeks before recall of the previous memory was assessed. We show that voluntary running causes forgetting of context fear memories in a neurogenesis-dependent manner, and that neurogenesis-induced forgetting is present in rats across behavioral tasks despite differences in complexity or reliance on spatial, context, or object memories. In addition, we asked whether stronger memories are less susceptible to forgetting by varying the strength of training. Even with a very strong training protocol in the CFC task, we still observed enhanced forgetting related to increased neurogenesis. These results suggest that forgetting due to neurogenesis is a conserved mechanism that aids in the clearance of memories.

**Significance Statement:** Recent evidence indicates that hippocampal neurogenesis mediates forgetting of older memories and enhances encoding of new memories free of proactive interference. This evidence comes from multiple rodent species, behavioral tasks, and methods of increasing neurogenesis. However, a recent paper by (Kodali et al. 2016) found that voluntary exercise-induced neurogenesis did not cause forgetting in the Morris Water Task in rats. The results call into question whether the phenomenon is a conserved function of neurogenesis across species. In the present study, we show that voluntary running causes robust forgetting in rats in a neurogenesis-dependent manner and that the effect is present across three different behavioral tasks, confirming the existence of the phenomenon in rats and adding to the growing evidence that forgetting is a conserved function of hippocampal neurogenesis.

## Introduction

Hippocampal neurogenesis plays a critical role in long-term memory, but its precise roles are not fully understood. Ablation of adult neurogenesis, or impairing the synaptic integration of adult-born neurons, impairs the acquisition of new memories (Snyder et al. 2005; Winocur et al. 2006; Arruda-Carvalho et al. 2014) and enhancement of neurogenesis improves subsequent learning (Van Praag et al. 1999). Only recently has significant attention been paid to the retrograde effects of adult neurogenesis on existing memories. Recent evidence has demonstrated that adult neurogenesis mediates forgetting of existing memories. First predicted computationally (Weisz and Argibay 2012), several experimental studies have confirmed that increased neurogenesis causes retrograde degradation of long-term memory in different hippocampus (HPC)-dependent tasks, across rodent species, and across the lifespan (Akers et al. 2014; Epp et al. 2016; Ishikawa et al. 2016; Gao et al. 2018; Cuartero et al. 2019). A beneficial outcome of this neurogenesis-induced reduction in memory retention is that it allows for the encoding of new memories free from proactive interference caused by older memories (Epp et al. 2016). The presence of this phenomenon in three different rodent species (mouse, degu, guinea pig) suggests that it is a conserved mechanism for ameliorating proactive memory interference.

Contrary to computational models (Weisz and Argibay 2012) and experiments using mice (Akers et al. 2014; Epp et al. 2016; Gao et al. 2018), a recent study using rats found that increasing adult neurogenesis did not cause forgetting of a spatial memory in the Morris Water Task (MWT) (Kodali et al. 2016). These authors trained rats in the MWT and increased neurogenesis using voluntary wheel running. Despite exercise causing a ∼2 fold increase in the number of new neurons in the DG, there was no difference in retention of the original platform location, indicating that increasing neurogenesis did not impair retention of a previously acquired spatial memory. On this basis, the authors called into question whether neurogenesis-induced forgetting is present in rats and thus, whether the phenomenon is conserved across species.

To date, neurogenesis-induced forgetting in the rat has not been assessed in any other behavioral tasks. However, neurogenesis-induced forgetting in mice has been demonstrated in the MWT (Akers et al. 2014; Epp et al. 2016), contextual fear conditioning (CFC) (Akers et al. 2014; Ishikawa et al. 2016), the Barnes maze (Cuartero et al. 2019; Akers et al. 2014), inhibitory avoidance (Ishikawa et al. 2016), and odor-context paired associates learning (Epp et al. 2016). Thus, it is premature to conclude that neurogenesis-induced forgetting is absent in rats on the basis of a single behavioral assay.

Additionally, there were major differences between the MWT training protocols used by (Kodali et al. 2016) and (Epp et al. 2016). (Kodali et al. 2016) conducted a greater number of days of MWT training and used longer trials. Overtraining of the MWT is known to increase mossy fiber plasticity (Ramírez-Amaya et al. 1999) and increased complexity of CA3 dendrites (Gómez-Padilla and Bello-Medina 2019) This could possibly leading to a more robust circuit supporting the memory. Stronger training protocols can make memory more resistant to neurogenesis-induced forgetting (Akers et al. 2014) and even to HPC lesions (Lehmann et al. 2009; Sutherland, Sparks, and Lehmann 2010; Lehmann and McNamara 2011). Hence, the null findings in rats (Kodali et al. 2016) could be a result of overtraining and a viable comparison would require the use of an equivalent training protocol to that used in the mouse studies (Akers et al. 2014; Epp et al. 2016).

Here, we investigated the existence of neurogenesis-induced forgetting in the rat. Using voluntary exercise as a reliable means of increasing neurogenesis, we assessed multiple HPC-dependent behavioral tasks and also investigated different strengths of memory training. We provide robust evidence that neurogenesis-induced forgetting is present in the rat. Thus, this phenomenon is not unique to mice and may be an evolutionarily conserved mechanism for managing memory interference in the HPC.

## Materials and Methods

### Subjects

A total of 152 male adult Long Evans rats (Charles River Laboratories, Kingston, NY, USA) in total were used in the experiments. Ninety-six were used in the CFC experiments, 28 were used in the MWT experiment, and 28 were used in paired associates learning (PAL). Rats were pair housed in standard cages on a 12:12-h lighting cycle and had *ad libitum* access to standard rat chow and water except in the case of rats used in the PAL experiment, which underwent food restriction to reduce their body weight to 90% of their free feeding weight. All experiments and procedures were approved by institutional Animal Care Committees and conformed to institutional and national ethical standards.

### Manipulation of Neurogenesis

Voluntary wheel running was used as a manipulation because it causes a reliable increase in neurogenesis (Van Praag et al., 1999b; Wojtowicz et al., 2008; Speisman et al., 2012; Akers et al., 2014; Epp et al., 2016; Kodali et al., 2016) and although running has other effects on the brain, the forgetting effect has been shown to occur specifically due to increases in neurogenesis (Epp et al., 2016). Half of the rats in each experiment were provided with continuous access to running wheels in their home cages (Lafayette, Scurry Rat Activity Wheel) in specialized cages in which rats were pair-housed. The amount of running was monitored in a subset of cages with odometers. Sedentary control rats remained pair-housed in standard caging.

To determine whether any changes in memory were related to neurogenesis or some other effect of running, we administered temozolamide (TMZ; Biosynth Carbosynth, San Diego, CA) to reduce neurogenesis. Rats were injected with 25 mg/kg TMZ or vehicle (10% DMSO in 0.9% Saline). Injections were given on 3 consecutive days per week followed by 4 days with no injection. Half of the rats from each treatment condition were sedentary while the other half were given running wheels. Using this design, the administration of TMZ in the runners is expected to block the increases in neurogenesis (Cuartero et al. 2019).

### Contextual Fear Conditioning

Fear conditioning was conducted in sound attenuated chambers with shocks delivered from a shock generator to a grated floor and freezing monitored via proprietary software (Med Associates Inc., Fairfax, VT). Rats were trained with two different protocols: a weak training protocol or a strong training protocol. Weak training (Figure 1A, 2A) involved the delivery of 3 shocks (1 mA, 2s) during a single 8 min session. Rats were given 2 min to explore the chamber prior to the delivery of the first shock with subsequent shocks spaced 1 min apart. Following the third shock, the rats remained in the chamber for 1 min before being removed to their homecage. The strong training protocol (Figure 2B) involved the repetition of the previously described weak training protocol over 3 consecutive days resulting in a total of 9 shocks. Following manipulations of neurogenesis, rats were returned to the fear conditioning chambers for retention testing in which freezing was monitored over a 5 min session and no shocks were delivered.

**Figure 1.**
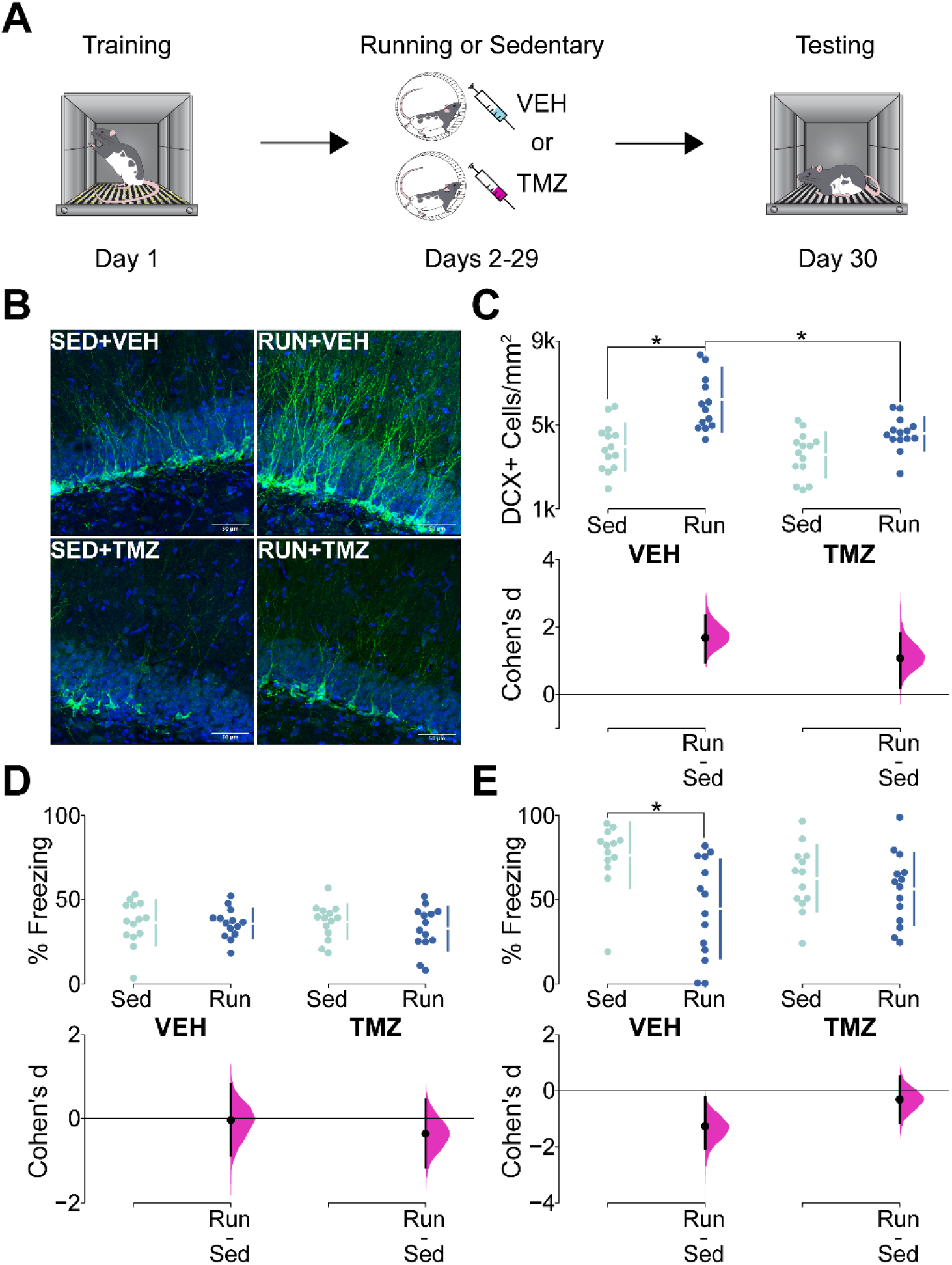
**A)** A Schematic of the behavior apparatus and experimental timeline used in contextual fear conditioning. **B)** Representative photomicrographs of DCX+ cells (green) in the DG (blue) of rats in the sedentary + vehicle (SED+VEH), runner + vehicle (RUN+VEH), sedentary + TMZ (SED+TMZ), and runner + TMZ (RUN+TMZ) groups. **C)** DCX+ cells/mm^2^ and effect sizes after sedentary control or running and vehicle or TMZ treatment. Upper portion of the plot shows individual data points in swarm plots with means (±SD; gapped vertical line). Lower portion of the plot shows effect size (Cohen’s d) as a black circle with vertical lines and gaussian distributions for the bootstrap 95% CI. Neurogenesis was substantially increased by running, and conversely, was decreased by TMZ. Moreover, this effect of TMZ was most striking in the running groups, with the RUN+TMZ groups exhibiting unchanged neurogenesis relative to the SED+TMZ group, but reduced neurogenesis relative to the RUN+VEH group. **D)** % Freezing and effect sizes in contextual fear conditioning prior to manipulation of neurogenesis. Upper portion of the plot shows individual data points in swarm plots with means (±SD; gapped vertical line). Lower portion of the plot shows effect size (Cohen’s d) as a black circle with vertical lines and gaussian distributions for the bootstrap 95% CI. All 4 groups exhibited equivalent levels of freezing during conditioning. **E)** % Freezing and effect sizes during CFC testing after manipulation of neurogenesis. Upper portion of the plot shows individual data points in swarm plots with means (±SD; gapped vertical line). Lower portion of the plot shows effect size (Cohen’s d) as a black circle with vertical lines and gaussian distributions for the bootstrap 95% CI. Freezing was significantly reduced in the RUN+VEH group relative to the SED+VEH group with no other significant differences being present. The results show that running causes forgetting relative to sedentary control, and that this effect is blocked by TMZ, indicating that the forgetting effect is dependent on increases in neurogenesis.

**Figure 2.**
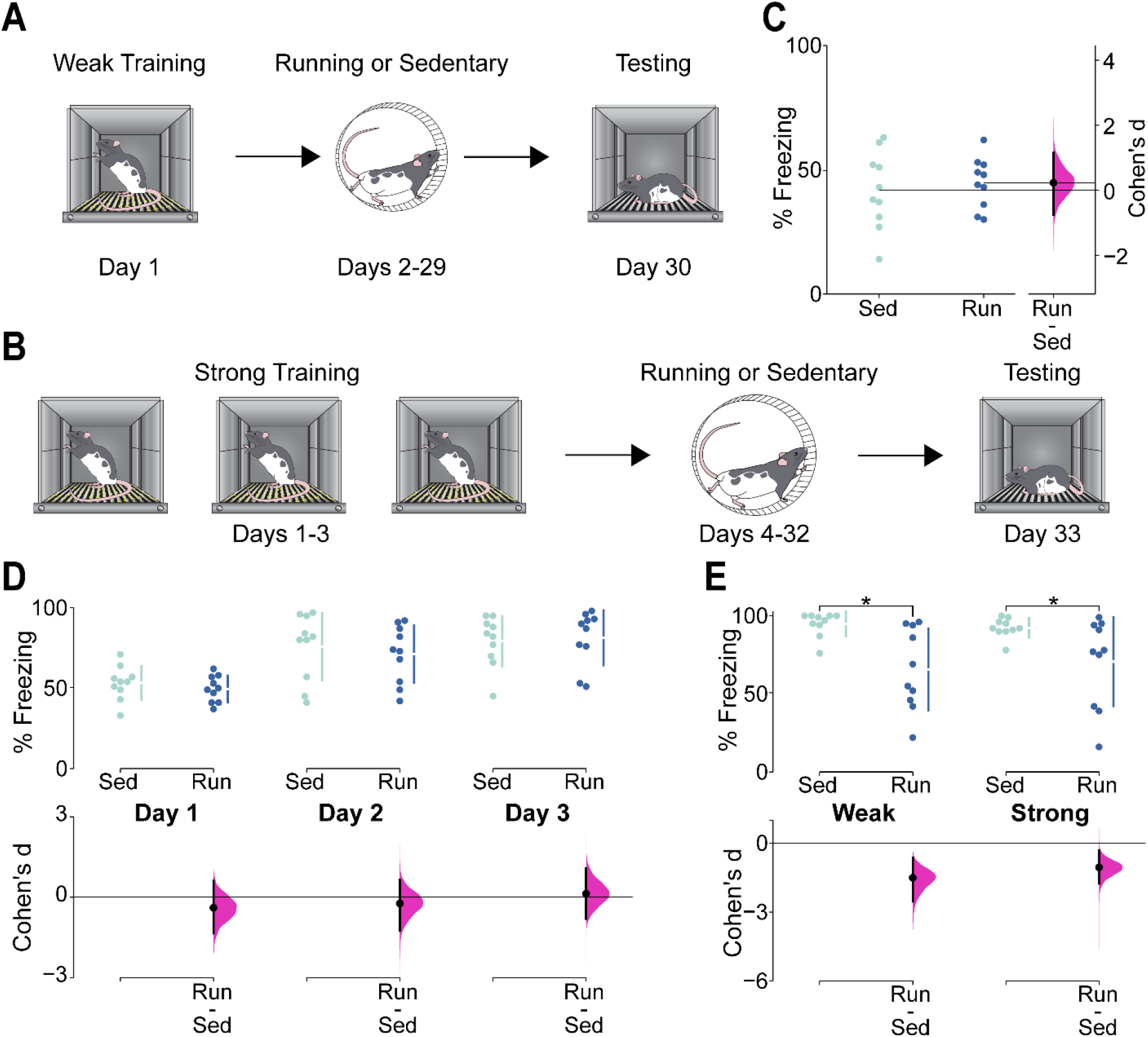
**A)** A schematic representation of the behavior and experimental timeline in the contextual fear conditioning weak training condition. **B)** A schematic representation of the behavior and experimental timeline in the contextual fear conditioning strong training condition. **C)** % Freezing during CFC training and effect size in the weak training condition for runners (Run) and sedentary controls (Sed). Data are shown as individual data points with mean (horizontal lines) and % Freezing scale on the left Y-axis. Effect size (Cohen’s d) is shown as a black circle with vertical lines and gaussian distribution for the bootstrap 95% CI and Cohen’s d scale on the right y-axis. Both groups froze the same amount during conditioning. **D)** % freezing during CFC training and effect sizes in the strong training condition. Upper portion of the plot shows individual data points in swarm plots with means (±SD; gapped vertical line). Lower portion of the plot shows effect size (Cohen’s d) as a black circle with vertical lines and gaussian distributions for the bootstrap 95% CI. Both groups froze the same amount during conditioning and percent freezing increased in both groups over successive days of conditioning. **E)** % Freezing in the weak training condition (Weak) and strong training condition (Strong) and effect sizes following either running or sedentary control. Upper portion of the plot shows individual data points in swarm plots with means (±SD; gapped vertical line). Lower portion of the plot shows effect size (Cohen’s d) as a black circle with vertical lines and gaussian distributions for the bootstrap 95% CI. Runners in both conditions showed large reductions in freezing, indicating that increasing neurogenesis caused forgetting, although the magnitude of the effect was somewhat lower in the strong training condition than in the weak training condition.

### Morris Water Task

MWT training (Figure 3A) was conducted in a pool (180 cm in diameter) filled with 21°C water and made opaque using non-toxic white tempera paint. A Plexiglas escape platform was submerged 2 cm below the surface of the water. Behavior was monitored using an overhead camera and Any-Maze tracking software (Stoelting Co., Wood Lane, IL). During initial training, rats were placed in one of the non-platform quadrants of the pool and allowed to swim until they located the platform, which was always located in the NW quadrant, or for a maximum of 60 s after which rats were placed on the platform for 15 s. We took several measures of rats’ swim patterns which included escape latency (s), distance (cm), and time inside the platform quadrant (s). Twenty-four hours after the final training session and prior to voluntary exercise, rats were administered a probe trial in which they were placed in the SW quadrant of the pool with the platform removed and were allowed to swim for 60 s to assess spatial preferences for the platform location.

**Figure 3.**
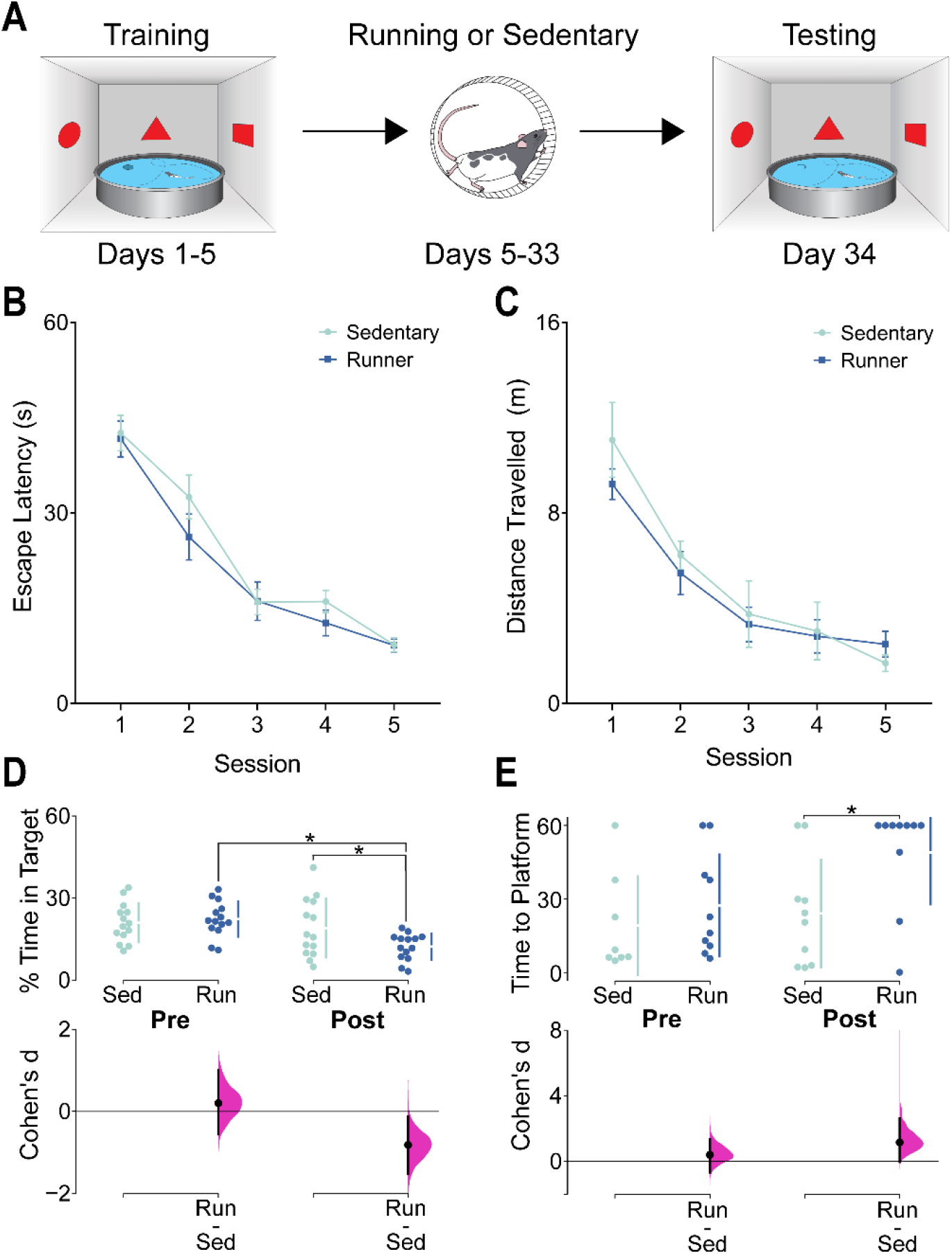
**A)** A schematic representation of behavior in the MWT and the timeline of behavior training and elevation of neurogenesis. **B)** Mean (±SEM) Escape Latency during MWT training. Both groups showed equivalent reductions in latency as training progressed. **C)** Mean (±SEM) Distance Travelled to the platform during MWT training. Both groups showed equivalent decreases in distance travelled as training progressed. **D)** % time in the platform zone (% Time in Target) and effect sizes during pre-running (Pre) and post-running (Post) probe trials for runners (Run) and sedentary controls (Sed). Upper portion of the plot shows individual data points in swarm plots with means (±SD; gapped vertical line). Lower portion of the plot shows effect size (Cohen’s d) as a black circle with vertical lines and gaussian distributions for the bootstrap 95% CI. Prior to manipulation of neurogenesis, runners and sedentary controls spent an equivalent percentage of time in the platform zone, but after increasing neurogenesis in the running condition, runners spent significantly less time in the platform zone relative to sedentary controls, indicating that running caused forgetting of the platform location. **E)** Latency to the first platform zone crossing (Time to Platform) and effects sizes during pre-running and post-running probe trials. Upper portion of the plot shows individual data points in swarm plots with means (±SD; gapped vertical line). Lower portion of the plot shows effect size (Cohen’s d) as a black circle with vertical lines and gaussian distributions for the bootstrap 95% CI. After manipulation of neurogenesis, runners took significantly longer to first cross the platform zone than controls, demonstrating that the diminished memory in runners was not an artifact of an altered search strategy.

Following neurogenesis manipulation, rats’ spatial memory was tested by returning them to the pool for a probe trial in which the platform was removed. Time spent in the platform quadrant was taken as a measure of memory for the original platform location. In addition, we quantified the latency to rats’ first crossing of the platform zone in order to rule out potential differences in the search strategies of controls and runners.

### Paired Associates Learning

#### Touchscreen Apparatus

PAL procedures were conducted within eight touchscreen-equipped operant conditioning chambers (Lafayette Instruments, Lafayette, IN, USA; An illustration of the interior of a chamber is shown in Figure 4A). Each chamber was contained within a sound-attenuating box and a fan for background noise and air circulation. A live video feed of animal activity was acquired through a camera mounted within the box above the operant chamber. An interchangeable mask with 3 rectangular windows of equal size was placed flush against the touchscreen and a spring-loaded shelf was located just below these windows, requiring animals to stand when making a response.

**Figure 4.**
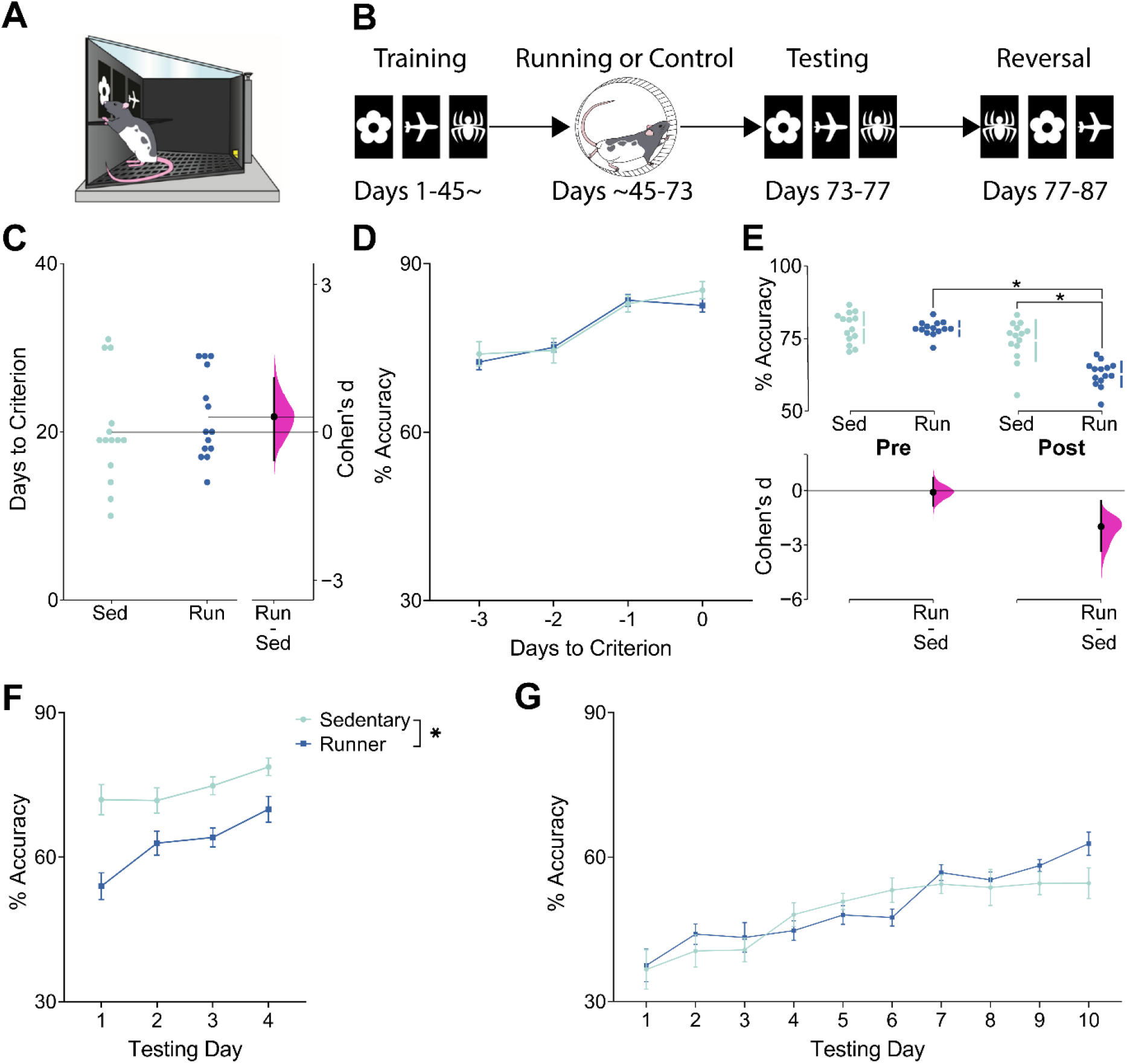
**A)** A schematic depicting the touchscreen apparatus used for PAL training and testing. **B)** A timeline depicting the sequence of experiments in PAL. **C)** Number of days to criterion for runners (Run) and sedentary controls (Sed) and effect size. Data are shown as individual data points with mean (horizontal lines) and # Days to Criterion scale on the left Y-axis. Effect size (Cohen’s d) is shown as a black circle with vertical lines and gaussian distribution for the bootstrap 95% CI and Cohen’s d scale on the right y-axis.. Both groups reached the initial criterion in PAL at the same rate before elevation of neurogenesis. **D)** Mean (±SEM) % Accuracy during the final 4 days of PAL training before elevation of neurogenesis. Both groups performed with the same accuracy before elevation of neurogenesis. **E)** % Accuracy in PAL training before (Pre) and after (Post) elevation of neurogenesis and effect sizes (% Accuracy is computed as an average across 4 days of testing for each swarm plot). Upper portion of the plot shows individual data points in swarm plots with means (±SD; gapped vertical line). Lower portion of the plot shows effect size (Cohen’s d) as a black circle with vertical lines and gaussian distributions for the bootstrap 95% CI. Both groups performed with the same % Accuracy before elevation of neurogenesis, but runners performed with significantly reduced accuracy after elevation of neurogenesis, indicating that they had forgotten the correct image-location pairings. **F)** Mean (±SEM) percent accuracy in PAL following either running or sedentary control. Runners performed with significantly and persistently reduced accuracy relative to sedentary controls over 4 days of testing. **G)** Mean (±SEM) % Accuracy in PAL during reversal learning. Both groups performed with the same accuracy during reversal learning, indicating that running did not enhance accuracy during PAL reversal learning.

#### Touchscreen Habituation and Pretraining

Animals were handled for at least 5 days before touchscreen habituation began. The first day of habituation involved acclimatizing the animals to the touchscreen room. Animals were transferred from the housing room to the touchscreen room and were given 5 reward pellets (Dustless Precision Pellets, 45 mg, Rodent Purified Diet; BioServ, NJ, USA) before being left undisturbed for 1 hour. During this period, all equipment was on and the lights were dimmed. For two additional days, animals were placed in the touchscreen chambers for 30 min and given 5 reward pellets. On all subsequent days of training or testing, animals were acclimatized in the touchscreen room for 15-20 min before going into the chambers.

Pretraining consisted of 4 progressive phases for training the animals to interact with the touchscreen display and receive rewards. 1) Initial Touch: One of the response windows was illuminated pseudorandomly for 30 s and 3 reward pellets were delivered if the rat correctly touched the illuminated window during this period or only 1 reward pellet if the illuminated window was not touched. Trials were interspersed with a 20 s intertrial period. Criterion for Initial Touch was completion of 100 trials in 1 h. 2) Must Touch: Identical to Initial Touch, except the animals received only 1 reward pellet and only for correct responses. The criterion for Must Touch training was 100 trials in 1 h. 3) Must Initiate: Identical to Must Touch except that the animal was required to initiate each trial by nose poking in the reward port. Criterion for the Must Initiate phase was 100 trials in 1 h. 4) Punish Incorrect: Identical to Must Initiate except that incorrect touches were punished with a 5-s time out and a correction trial. Correction trials are identical to the previous presentation and repeat until successfully completed. The criterion for Punish Incorrect was 100 trials in 1 h, with greater than 80% correct, with accuracy calculated for the initial presentation only.

#### PAL Training

PAL required the animal to differentiate between two different images presented simultaneously in 2 of the 3 response windows pseudorandomly. Each image is correct only when paired with its respective location. Negative images of a flower, airplane, and spider were used as stimuli. The flower is always correct in the left position, the airplane in the centre position, and the spider in the right position. Correct responses are rewarded and punished in the same manner in the Punish Incorrect phase. Animals were trained to a criterion of 90 correct selection trials, completed within 1 h, with greater than 80% accuracy, for two consecutive days. Correction trials were not included in the number of selection trials completed or the calculation of accuracy scores.

A schematic of the experimental timeline used in the PAL experiment is shown in Figure 4B. After animals reached criterion in PAL, cage-mates were randomly assigned to a running wheel cage or maintained in their standard home cage. Each pair of cage-mates assigned to the running wheel condition were yoked to another pair that would be assigned to the sedentary condition in order to maintain counterbalancing of the number of pre-training days between experimental conditions. At the end of the neurogenesis protocol all animals were food restricted to 90% of free feeding and placed in new standard home cages. Animals were given 1 day to acclimatize to the transfer procedure.

#### PAL Testing and Reversal Learning

Following the neurogenesis protocol animals PAL retention was tested with 4 sessions of PAL. After PAL animals were moved to reversal regardless of performance and tested daily, seven days a week, for 10 days. Reversal was identical to PAL except that the correct stimuli/location pairings were changed so that the airplane was correct in the left position, the spider in the centre position, and the flower in the right position.

### Perfusions and Immunohistochemistry

After the conclusion of behavioral testing, rats were deeply anaesthetized and perfused intracardially. Brains were stored for 24 hours in 4% formaldehyde at 4° C before being transferred to a cryoprotectant solution composed of 30% sucrose/0.1% sodium azide. Tissue was sectioned using a cryostat (Leica) and 12 series of 40 µm thick tissue sections were collected into an antifreeze solution composed of PBS buffered glycerol and ethylene glycol for storage at −20°C. Immunohistochemistry was conducted to label DCX-positive neurons in the dentate gyrus. Tissue was washed 3 times in 0.1M PBS before being transferred to a primary antibody solution containing 0.1M PBS with 3% Trition-X, 3% Donkey Serum, and a 1:200 dilution of rabbit anti-DCX primary antibody (Product #4604S, Cell Signalling Technology, Danvers, MA). Tissue was incubated in the primary antibody solution at room temperature for 48 h before being washed 3 times in 0.1M PBS and transferred to a secondary antibody solution containing 0.1M PBS and a 1:500 dilution of donkey anti-rabbit antibody (Dylight 488). Sections were then transferred to a solution containing 0.1M PBS and a 1:2000 dilution of DAPI before being mounted to glass slides and coverslipped with polyvinyl alcohol/DABCO mounting medium.

### Quantification of DCX

Quantification of DCX+ granule cells was performed throughout the entire rostral-caudal extent of the DG using a fluorescence microscope (Olympus FV3000) with 60x NA 1.35 oil objective. Cells were counted in the granule cell layer and the subgranular zone (defined as the 50 µm zone adjacent to the granule cell layer). The area of the DG in every section was traced in order to estimate DG volume. DCX+ cell counts were normalized to DG volume to control for differences in DG volume between subjects. Quantification was performed blind to treatment conditions.

### Experimental Design and Statistical Analysis

We used a combination of traditional hypothesis testing and estimation statistics, which relies on quantitative judgements of effect sizes rather than significance thresholds (Ho et al. 2019). Continuous data such as learning curves were typically analyzed and expressed as traditional plots with standard parametric statistics, whereas post-running group differences were evaluated using combined estimation statistics and hypothesis testing. In such cases, data were represented as estimation plots with significant p-values obtained from standard hypothesis testing and displayed on the plots. Data displayed includes individual subjects as swarm plots in addition to showing group means and standard deviation (gapped vertical line). Effect sizes are shown in separate plots as bootstrap 95% confidence intervals and were calculated as Cohen’s d. Estimation plots and statistics were generated using the Estimation Statistics website (estimationstats.com). Standard hypothesis testing analyses were performed in Prism 9.

## Results

### Voluntary Running Causes Forgetting of Contextual Fear Conditioning in a Neurogenesis-Dependent Manner

As expected based on previous literature, we observed significant main effects of running (F(1,52) ≤ 27.79, p < 0.0001)) and of TMZ (F(1,52) ≤ 10.45, p < 0.0021) in the number of DCX+ cells in the DG (Representative photomicrographs shown in Figure 1B). We also found a significant running × treatment interaction effect (F(1,52) = 4.06, p = 0.049). Tukey’s post hoc test demonstrated that there was a significant increase in DCX+ cells in vehicle treated runners compared to vehicle treated sedentary mice (*p < 0*.*0001)*. However, there was no significant increase in DCX+ cells in the TMZ treated runners compared to the TMZ treated sedentary group (*p = 0*.*11)*. Importantly, treatment with TMZ did not cause a statistically significant reduction in the amount of running (RUN+VEH 5.43(±0.85) km/day/rat, RUN+TMZ 3.44(±0.52) km/day/rat; t(10) = 1.98, p = 0.076).

Prior to manipulation of neurogenesis with running/TMZ, all 4 groups exhibited equivalent levels of freezing during CFC acquisition and there were no significant main or interaction effects (Figure 1D: Fs(1,52) ≤ 0.53, ps ≥ 0.45). After manipulation of neurogenesis, we found a significant running by treatment interaction (Figure 1E. Significant interaction of Running × TMZ: F(1,50) = 4.14, p = 0.047). Post hoc tests indicated that the RUN+VEH group froze significantly less than the SED+VEH group (*p = 0*.*0039)*, whereas this effect of running on fear memory was absent in the TMZ treated mice (*p = 0*.*89)*.

### Increasing Neurogenesis Causes Forgetting of Contextual Fear Condition Regardless of Strength of Training

Having established that the forgetting effect in CFC is neurogenesis-dependent, we sought to determine whether this effect could also be modulated by the strength of CFC training. Runners and sedentary controls exhibited similar levels of freezing during conditioning in the weak training (Figure 2C: t(18) = 0.53, p =.60) and strong training (Figure 2D: F(1,54) = 0.25, p = 0.62), although both groups in the strong training condition exhibited increased levels of freezing over successive conditioning sessions (Figure 2D; Main effect of Day: F(2,54) = 18.81, p <0.0001). After voluntary running (running effects on DCX shown in Supplemental Materials Figure S1), runners froze significantly less than sedentary controls in both the weak and strong training conditions (Figure 2E. Significant main effect of group F(1,36) = 16.21. p = 0.0003), indicating weakened memory for the context-fear association in runners regardless of strength of training. There was no significant interaction effect when comparing freezing in the weak and strong training conditions (F(1,36) = 0.34, p = 0.56), but a raw comparison of the effect sizes reveals that the forgetting effect in the strong training condition was of ∼30% lower magnitude than in the weak training condition.

### Increasing Neurogenesis Causes Forgetting in the MWT

We next examined whether running induced neurogenesis altered memory retention using a spatial version of the MWT. Prior to increasing neurogenesis, runners and sedentary controls both acquired the MWT equally with equivalent decreases in escape latency (Figure 3B. Significant main effect of Training Day: F(4,26) = 75.16, p < 0.0001) and distance travelled to the platform (Figure 3C. Significant main effect of Training Day: F(4,16) = 25.17, p < 0.0001). Probe trials conducted before and after elevation of neurogenesis showed that runners exhibited a reduction in memory following 4 weeks of running but sedentary mice showed no significant change (Figure 3D. Significant interaction of Group × Probe Trial: F(1,52) = 4.07, p = 0.049). After elevation of neurogenesis, runners spent significantly less time in the target zone they did during the pre-running probe trial *(p = 0*.*005)* and less than sedentary controls in the post-running probe trial *(p = 0*.*042)*. Additionally, runners had a significantly longer latency to their first platform zone crossing in the post-running probe trial showing that the memory impairment in runners was not due to any enhanced adaptation in platform search strategy (Figure 3E. Significant main effect of Group: F(1,18) = 5.22, p = 0.03. Significant post-neurogenesis post hoc test; p = 0.03).

### Increasing Neurogenesis Causes Forgetting of PAL

As a final test of whether neurogenesis causes forgetting, we examined the effects of increased neurogenesis on performance in PAL. Prior to elevation of neurogenesis, both groups achieved criterion in the same amount of time (Figure 4C) and performed with nearly identical % Accuracy in PAL (Figure 4D,E). Analyses of pretraining Trial Completion and Response Latency are shown in the Supplemental Materials (Figure S2A-B. Figure S3A-C, Table S1). After 4 weeks of voluntary exercise or sedentary control, runners exhibited a large and significant reduction in % Accuracy (Figure 4E. Significant interaction of Neurogenesis × Test: F(1,26) = 27.10, p < 0.0001). Moreover, the performance deficit, relative to sedentary controls, was sustained over 4 days of testing (Figure 4D,F; Significant main effect of Neurogenesis: F(1,26) = 27.29, p < 0.0001), although both groups also significantly improved their performance over the 4 days of retention testing, indicating significant relearning of the task (Figure 4F; Significant main effect of Testing Day: F(3,78) = 9.49, p = 0.0002). The lack of a significant interaction of Group × Testing Day suggests that both runners and controls underwent significant improvements to % Accuracy (F(3,78) = 2.00, p = 0.12). All other analyses of post-neurogenesis PAL performance are included in the Supplemental Materials (Figure S2C-D, Figure S3D-F, Table S2). Briefly, there were no differences in Trial Completion or Response Latency prior to increasing neurogenesis, but after increasing neurogenesis, runners performed more correction trials (Figure S2D. Significant main effect of Group: F(1,26) = 27.16, p < 0.0001) and performed with reduced Incorrect Choice Latency (Figure S4E. Significant main effect of Group: F(1,26) = 6.43, p = 0.02). The altered Incorrect Choice Latency was not correlated with % Accuracy (r(26) = 0.04, p = 0.68) meaning that the reduction in % Accuracy was likely not the result of a change in response time and represents forgetting rather than a speed-accuracy trade-off effect.

### Increased Neurogenesis Does Not Improve Reversal Learning Accuracy, but Increases Behavioral Flexibility and Response Speed

We also sought to determine whether increased neurogenesis would facilitate reversal learning in PAL. There were no significant differences between runners and sedentary controls in % Accuracy (Figure 4F; F(1,26) = 0.45, p = 0.50), and no interaction of Group × Testing Day (F(9,234) = 1.40, p = 0.18), indicating that increased neurogenesis did not enhance accuracy during reversal learning. However, there was a main effect of Training Day (F(9,234) = 17.65, p < 0.0001) showing that both groups significantly improved their accuracy over successive testing days. Analyses of PAL reversal learning Trial Completion and Response Latency are shown in the Supplemental Materials (Figure S2E-F, Figure S3G-I, Table S3). Runners also performed with increased Selection Trials relative to sedentary controls (Figure S2E. Significant main effect of Neurogenesis: F(1,26) = 26.97, p < 0.0001) and reduced the number of Correction Trials they performed over successive days of reversal learning whereas sedentary controls did not (Figure S3F. Significant Interaction of Neurogenesis × Testing Day: F(9,234) = 2.00, p = 0.04). Finally, runners performed with decreased Correct Choice Latency (Figure S3G. Significant main effect of Neurogenesis: F(1,26) = 24.02, p < 0.0001), Incorrect Choice Latency (Figure S3H. Significant main effect of Neurogenesis: F(1,26) = 18.56, p < 0.0001), and Reward Collection Latency (Figure S3I. Significant main effect of Neurogenesis: F(1,26) = 5.50, p = 0.027). All other analyses of PAL reversal learning performance are shown in Tables 4-5.

## Discussion

In the present study, we investigated the effects of increasing hippocampal neurogenesis, via voluntary running, on memory retention in three different HPC-dependent memory tasks. We show here consistent evidence that voluntary running results in forgetting of previously acquired memories in rats in a neurogenesis-dependent manner and that the effect extends to 3 different HPC-dependent long-term memory tasks. Several previous reports using mice, guinea pigs, and degus have demonstrated that increasing neurogenesis causes forgetting of previously acquired hippocampal memories (Akers et al. 2014; Epp et al. 2016; Ishikawa et al. 2016; Gao et al. 2018; Cuartero et al. 2019). However, a recent report found a null result in rats, calling into question whether the neurogenesis-induced forgetting phenomenon is present in rats and, thus, whether it is an evolutionarily conserved mechanism (Kodali et al. 2016). Our current findings show that neurogenesis-induced forgetting is clearly present in rats.

The debate around this issue is of central importance to fields involving long-term memory research under both normal and pathological conditions. Forgetting is an essential complementary process to memory (Medina 2018), aiding in the balance between plasticity and the stability of memory circuits. Thus, understanding the mechanisms underlying forgetting, including neurogenesis, is critical to the broader understanding of long-term memory function.

Neurogenesis was substantially and similarly increased by voluntary exercise, replicating a large body of previous findings (Van Praag et al. 1999; Van Praag, Kempermann, and Gage 1999; Wojtowicz, Askew, and Winocur 2008; Speisman et al. 2012; Akers et al. 2014; Epp et al. 2016; Kodali et al. 2016). Additionally, the running-induced increase in neurogenesis was blocked by TMZ administration in a similar fashion to previous findings (Akers et al. 2014; Epp et al. 2016). Although both running and TMZ administration have effects outside of changes in neurogenesis, transgenic approaches to blocking neurogenesis have also yielded the same results (Akers et al. 2014).

We found that contextual fear memory was significantly impaired by voluntary exercise and that the effect could be blocked by treatment with TMZ to inhibit neurogenesis, replicating previous findings (Akers et al. 2014; Epp et al. 2016; Ishikawa et al. 2016; Cuartero et al. 2019). We found that both strong and weak training of the contextual fear memories resulted in neurogenesis-induced forgetting. Although the forgetting effect was of slightly lower magnitude following a strong training protocol, suggesting that more strongly trained memories may be somewhat more resilient to the effects of neurogenesis, both training protocols proved to be highly susceptible to the forgetting effect. This suggests that even highly salient memories are vulnerable to disruption by elevated neurogenesis.

In direct contrast to the results of (Kodali et al. 2016), we found that voluntary exercise caused forgetting of a previous platform location in the MWT, replicating previous findings that elevated neurogenesis causes retrograde impairments in spatial memory (Akers et al. 2014; Epp et al. 2016; Cuartero et al. 2019). The rats in the Kodali et al (2016) study were administered these sessions over a period of 8 days rather than 4 days. Previous research involving HPC lesions has shown that memory acquired in more distributed learning sessions can become independent of the HPC (Lehmann et al. 2009). Thus, the discrepancy here may relate to how MWT training sessions are distributed over time rather than the raw duration of training. Additionally, we measured the latency to the first platform zone crossing and found that runners were significantly impaired on this variable as well, indicating that our results did not come as a result of altered search strategy in the MWT.

Increasing neurogenesis also caused a large decrease in accuracy of the previously acquired PAL task. It has been shown that increasing neurogenesis disrupts odor-based PAL in mice (Epp et al. 2016), but the present results are the first published account of neurogenesis-induced forgetting in the touchscreen-based PAL task. Importantly, the touchscreen-based PAL task differs from both the MWT and contextual fear conditioning in that it requires extensive training over a period of weeks but, despite this, we again show the presence of neurogenesis-induced forgetting.

Increased neurogenesis did not enhance accuracy during reversal learning, counter to previous findings that enhancing neurogenesis alleviates proactive interference (Epp et al. 2016). This may have been related to the number of days of retention testing. Over the course of retention testing, animals in both groups showed improvements in performance consistent with relearning of the original image-location associations. Thus, reversal learning may have been impaired by interference from relearning during retention testing. However, runners performed significantly more trials than controls during reversal. Additionally, runners exhibited a greater reduction in the number of correction trials than controls over successive days of reversal learning. Runners also performed with lower latency than controls across all latency measures. Correction trials have previously been used as a measure of cognitive flexibility (Lins, Phillips, and Howland 2015; Lins and Howland 2016). The combined increase in selection trials, enhanced reduction in correction trials, and lower latencies can also be considered an indication of a general increase in task efficiency (Roebuck et al. 2018). In brief, runners performed PAL reversal learning faster and more flexibly than sedentary controls, if not more accurately.

Combined, these results show that neurogenesis-induced forgetting is a clear and large effect in rats across tasks. It is worth emphasizing, however, that these results are not in contradiction with previous studies showing the necessity of adult-born neurons for recall of previously-encoded long-term memories. If adult-born neurons are ablated after memory formation, recall of those memories is impaired presumably because the ablated new neurons had become integrated as necessary units in the memory trace (Arruda-Carvalho et al. 2011). Our present claim relates specifically to the retrograde effects on long-term memories of adult-born neurons that have been *newly generated after* the formation of a memory trace.

In conclusion, the present study sought to extensively test whether neurogenesis-induced forgetting is present in rats. Using 3 different behavioral tasks involving different types of HPC-dependent memory with differential complexity, strength of training, and sensory modality, we show that neurogenesis-induced forgetting is robustly present in rats across HPC-dependent memory, replicating a body of previous literature in other rodent species.

## Acknowledgements

Funding for this study was provided by an NSERC Discovery Grant (RGPIN-2018-05135) to JRE and an NSERC Discovery Grant and CIHR operating Grant to JGH. GAS received funding from NSERC, the Hotchkiss Brain Institute and the Cumming School of Medicine. AJR received funding from NSERC.

