## Supplemental Materials for "Adult neurogenesis mediates forgetting of multiple types of memory in the rat"

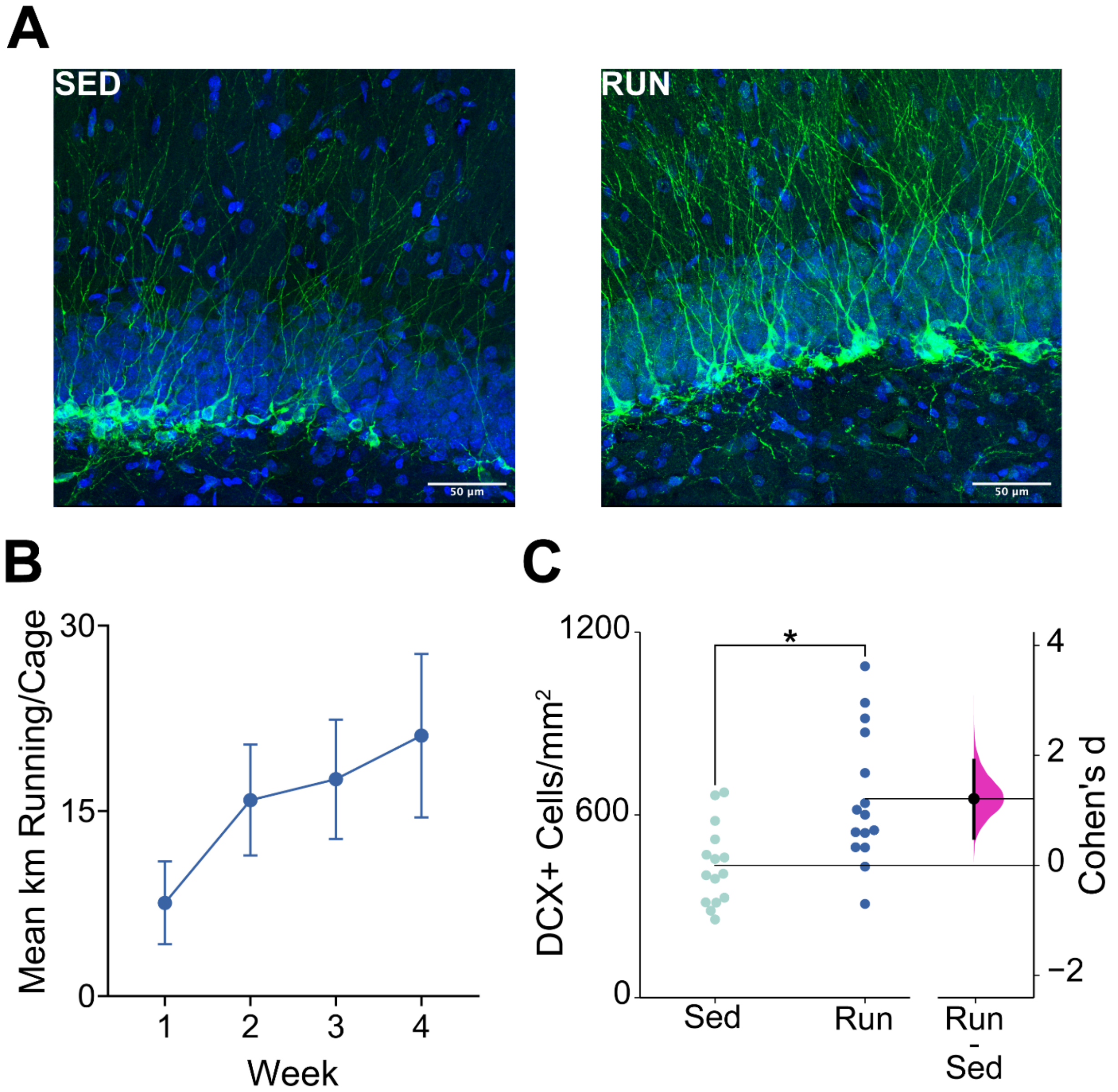


**Figure S1. A)** Representative photomicrographs of DCX-positive cells in the DG of sedentary controls (Sed) and runners (Run). **B)** Mean (±SEM) kms of running (per cage) completed by runners during the 4 weeks of voluntary running. **C)** DCX+ cells/mm$^2$. Data are shown as individual data points with mean (horizontal lines) and DCX+ cells/mm$^2$ scale on the left Y-axis. Effect size (Cohen's d) is shown as a black circle with vertical lines and gaussian distribution for the bootstrap 95% CI and Cohen's d scale on the right y-axis. Running caused a significant increase in the number of DCX-positive cells, indicating that neurogenesis was robustly increased.

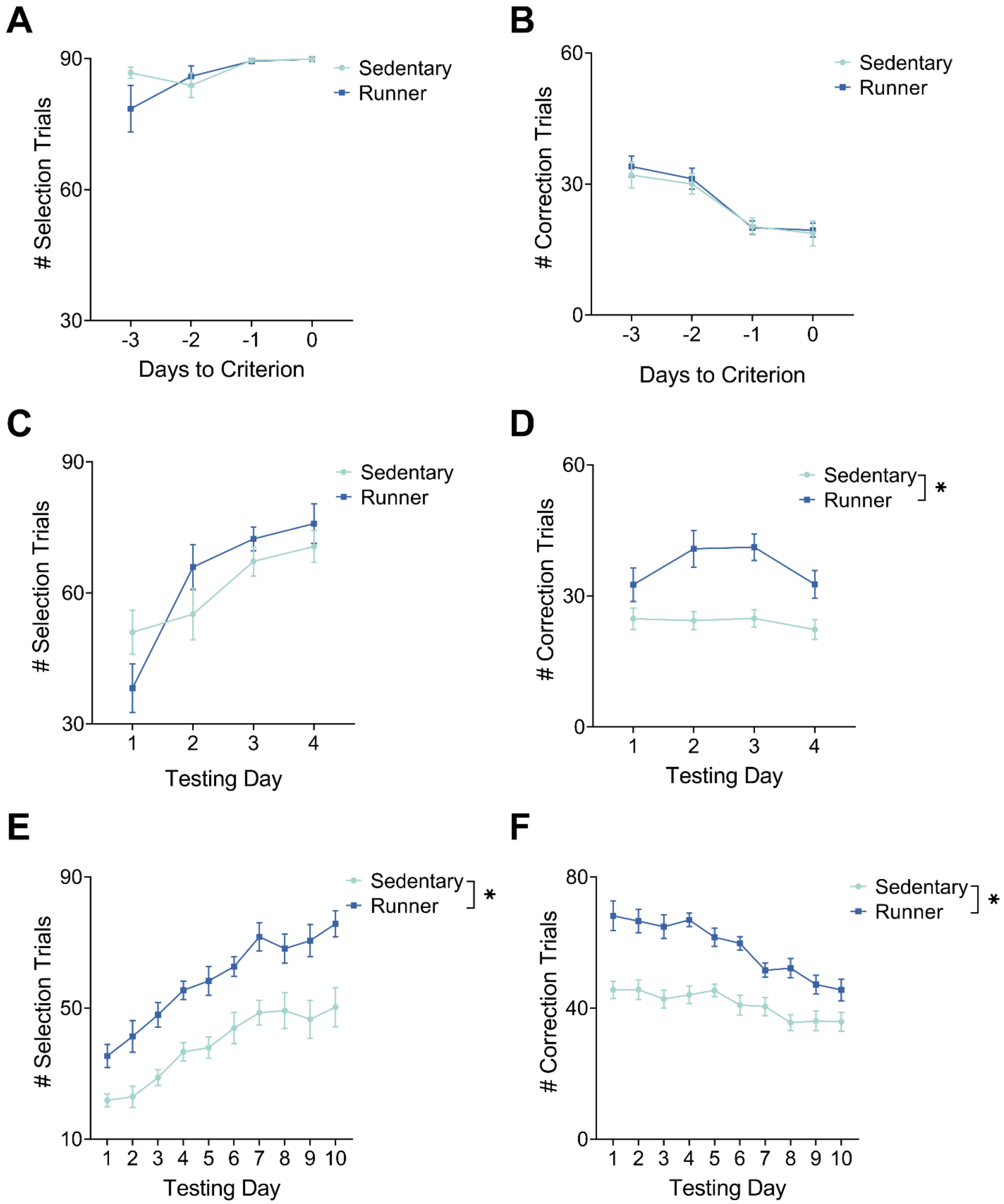
A
### Selection Trials
Sedentary
Runner
Days to Criterion
B
### Correction Trials
Sedentary
Runner
Days to Criterion
C
### Selection Trials
Sedentary
Runner
Testing Day
D
### Correction Trials
Sedentary
Runner
*
Testing Day
E
### Selection Trials
Sedentary
Runner
*
Testing Day
F
### Correction Trials
Sedentary
Runner
*
Testing Day

**Figure S2. A)** Mean (±SEM) number of selection trials performed by runners and sedentary controls during pretraining in PAL. Both groups performed roughly the same number of selection trials. **B)** Mean (±SEM) number of correction trials performed by runners and sedentary controls during pretraining in PAL. Both groups performed an equivalent number of correction trials during pretraining. **C)** Mean (±SEM) number of selection trials performed by runners and sedentary controls during testing in PAL after elevation of neurogenesis. Runners and controls performed an equivalent number of selection trials. **D)** Mean (±SEM) number of correction trials performed by runners and sedentary controls during testing in PAL after elevation of neurogenesis. Runners performed significantly more correction trials than controls. **E)** Mean (±SEM) number of selection trials performed by runners and sedentary controls during reversal learning. Runners performed significantly more selection trials than sedentary controls, indicating enhanced motivation to perform the PAL task in the presence of altered image-location associations. **F)** Mean (±SEM) number of correction trials performed by runners and sedentary controls during reversal learning. Runners performed significantly more corrections trials. However, runners also reduced the number of correction trials they performed over successive days in contrast to controls, indicating that running caused an enhancement of cognitive flexibility during reversal learning.

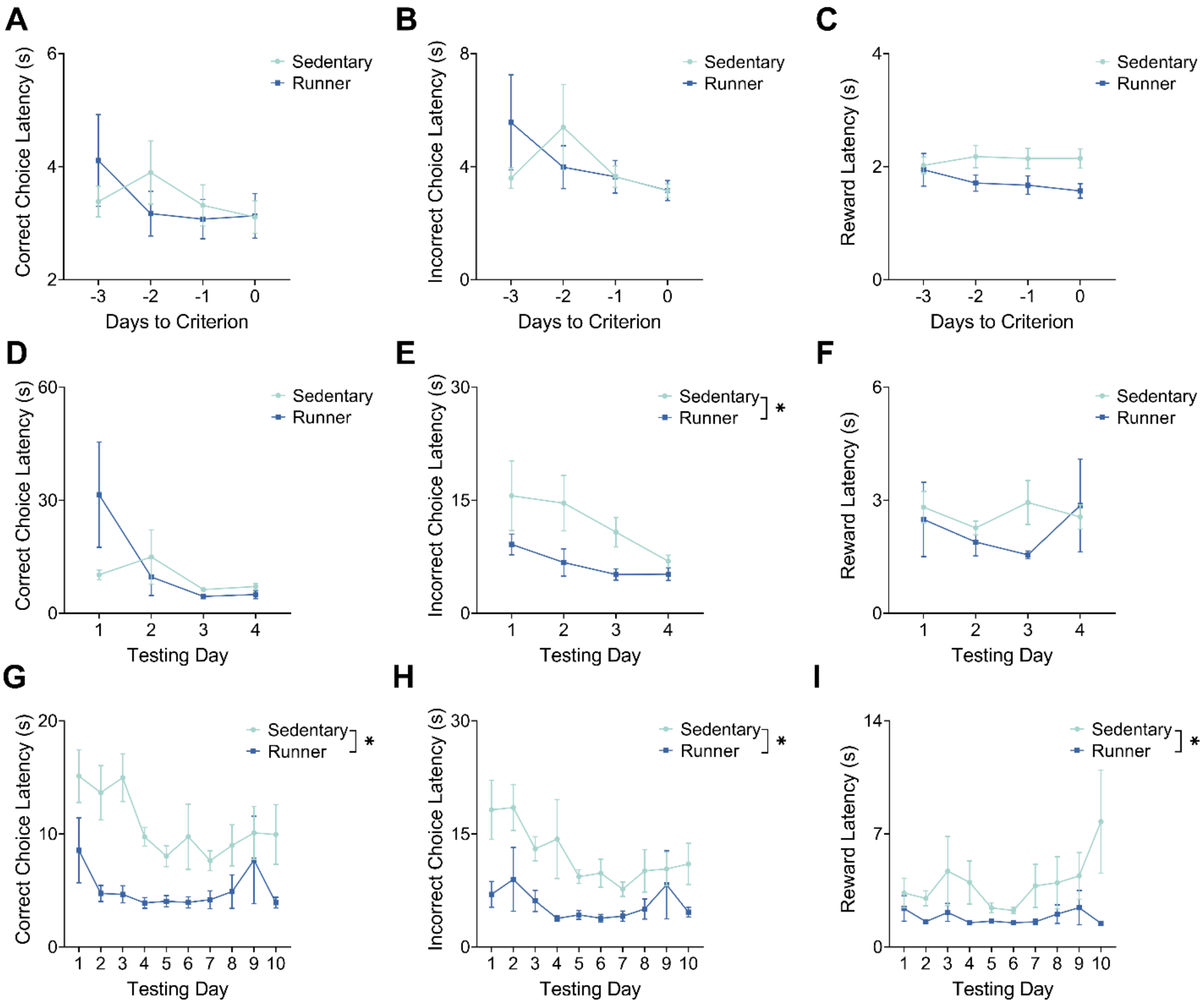


**Figure S3. A)** Mean (±SEM) correct choice latency in runners and sedentary controls prior to elevation of neurogenesis. Both groups performed with equivalent correct choice latency. **B)** Mean (±SEM) incorrect choice latency in runners and sedentary controls prior to elevation of neurogenesis. Both groups performed with equivalent incorrect choice latency. **C)** Mean (±SEM) reward latency in runners in sedentary controls prior to elevation of neurogenesis. Both groups performed with equivalent reward latency. **D)** Mean (±SEM) correct choice latency in runners and sedentary controls after elevation of neurogenesis. Both groups performed with equivalent correct choice latency. **E)** Mean (±SEM) incorrect choice latency in runners and sedentary controls after elevation of neurogenesis. Runners performed with significantly lower incorrect choice latency than sedentary controls, but incorrect choice latency was not correlated with performance. **F)** Mean (±SEM) reward latency in runners and sedentary controls after elevation of neurogenesis. Both groups performed with equivalent reward latency. **G)** Mean (±SEM)

correct choice latency in runners and sedentary controls during reversal learning. Runners performed with significantly reduced correct choice latency relative to sedentary controls. **H)** Mean (±SEM) incorrect choice latency in runners and sedentary controls during reversal learning. Runners performed with significantly reduced incorrect choice latency. **I)** Mean (±SEM) reward latency in runners and sedentary controls during reversal learning. Runners performed with significantly reduced reward latency.

**Table S1**

***Complete Statistical Outputs for the Analysis of PAL Pretraining.***

| | Comparison | F | p |
|---|---|---|---|
| *Accuracy* | | | |
| | *Main Effect: Neurogenesis* | *F(1,26) = 0.19* | *.66* |
| | ***Main Effect: Training Day*** | ***F(3,78) = 45.8*** | ***<.0001*** |
| | *Neurogenesis X Training Day* | *F(3,78) = 0.98* | *.41* |
| *Correct Response Latency* | | | |
| | *Main Effect: Neurogenesis* | *F(1,26) = 0.01* | *.92* |
| | *Main Effect: Training Day* | *F(3,78) = 2.48* | *.10* |
| | *Neurogenesis X Training Day* | *F(3,78) = 2.64* | *.06* |
| *Incorrect Response Latency* | | | |
| | *Main Effect: Neurogenesis* | *F(1,26) = 0.02* | *.88* |
| | *Main Effect: Training Day* | *F(3,78) = 2.15* | *.12* |
| | *Neurogenesis X Training Day* | *F(3,78) = 1.89* | *.14* |
| *Reward Collection Latency* | | | |
| | *Main Effect: Neurogenesis* | *F(1,26) = 3.07* | *.09* |
| | *Main Effect: Training Day* | *F(3,78) = 0.56* | *.57* |
| | *Neurogenesis X Training Day* | *F(3,78) = 2.39* | *.08* |
| *Selection Trial Completion* | | | |
| | *Main Effect: Neurogenesis* | *F(1,26) = 0.61* | *.44* |
| | ***Main Effect: Training Day*** | ***F(3,78) = 5.37*** | ***.01*** |
| | *Neurogenesis X Training Day* | *F(3,78) = 2.20* | *.09* |
| *Correction trial Completion* | | | |
| | *Main Effect: Neurogenesis* | *F(1,26) = 0.14* | *.71* |
| | ***Main Effect: Training Day*** | ***F(3,78) = 30.85*** | ***<.0001*** |
| | *Neurogenesis X Training Day* | *F(3,78) = 0.13* | *.94* |

**Table S2**

***Complete Statistical Outputs for the Analysis of PAL Testing.***

| | *Comparison* | *F* | *p* |
|---|---|---|---|
| *Accuracy* | | | |
| | ***Main Effect: Neurogenesis*** | ***F(1,26) = 27.29*** | ***<.0001*** |
| | ***Main Effect: Testing Day*** | ***F(3,78) = 9.49*** | ***.0002*** |
| | *Neurogenesis X Testing Day* | *F(3,78) = 2.00* | *.12* |
| *Correct Response Latency* | | | |
| | *Main Effect: Neurogenesis* | *F(1,26) = 0.41* | *.52* |
| | *Main Effect: Testing Day* | *F(3,78) = 3.29* | *.0544* |
| | *Neurogenesis X Testing Day* | *F(3,78) = 2.41* | *.07* |
| *Incorrect Response Latency* | | | |
| | ***Main Effect: Neurogenesis*** | ***F(1,26) = 6.43*** | ***.0176*** |
| | ***Main Effect: Testing Day*** | ***F(3,78) = 3.46*** | ***.0395*** |
| | *Neurogenesis X Testing Day* | *F(3,78) = 0.76* | *.52* |
| *Reward Collection Latency* | | | |
| | *Main Effect: Neurogenesis* | *F(1,26) = 0.81* | *.3757* |
| | *Main Effect: Testing Day* | *F(3,78) = 0.49* | *.5946* |
| | *Neurogenesis X Testing Day* | *F(3,78) = 0.64* | *.5926* |
| *Selection Trial Completion* | | | |
| | *Main Effect: Neurogenesis* | *F(1,26) = 0.37* | *.5436* |
| | ***Main Effect: Testing Day*** | ***F(3,78) = 17.34*** | ***< .0001*** |
| | *Neurogenesis X Testing Day* | *F(1,78) = 2.49* | *.0642* |
| *Correction trial Completion* | | | |
| | ***Main Effect: Neurogenesis*** | ***F(1,26) = 27.16*** | ***<.0001*** |
| | *Main Effect: Testing Day* | *F(3,78) = 1.94* | *.1428* |
| | *Neurogenesis X Testing Day* | *F(3,78) = 1.19* | *.3194* |

**Table S3.**

***Complete Statistical Outputs for the Analysis of PAL Reversal Learning.***

| | *Comparison* | *F* | *p* |
|---|---|---|---|
| *Accuracy* | | | |
| | *Main Effect: Neurogenesis* | *F(1,26) = 0.45* | *.5* |
| | ***Main Effect: Training Day*** | ***F(9,234) = 17.65*** | ***<.0001*** |
| | *Neurogenesis X Training Day* | *F(9,234) = 1.40* | *.1892* |
| *Correct Response Latency* | | | |
| | ***Main Effect: Neurogenesis*** | ***F(1,26) = 24.02*** | ***<.0001*** |
| | *Main Effect: Training Day* | *F(9,234) = 2.37* | *.051* |
| | *Neurogenesis X Training Day* | *F(9,234) = 0.96* | *.47* |
| *Incorrect Response Latency* | | | |
| | ***Main Effect: Neurogenesis*** | ***F(1,26) = 18.56*** | ***.0002*** |
| | *Main Effect: Training Day* | *F(9,234) = 2.28* | *.0622* |
| | *Neurogenesis X Training Day* | *F(9,234) = 0.78* | *.6378* |
| *Reward Collection Latency* | | | |
| | ***Main Effect: Neurogenesis*** | ***F(1,26) = 5.496*** | ***.027*** |
| | *Main Effect: Training Day* | *F(9,234) = 1.27* | *.2921* |
| | *Neurogenesis X Training Day* | *F(9,234) = 1.272* | *.2529* |
| *Selection Trial Completion* | | | |
| | ***Main Effect: Neurogenesis*** | ***F(1,26) = 26.97*** | ***<.0001*** |
| | ***Main Effect: Training Day*** | ***F(9,234) = 28.44*** | ***<.0001*** |
| | *Neurogenesis X Training Day* | *F(9,234) = 0.55* | *.833* |
| *Correction trial Completion* | | | |
| | ***Main Effect: Neurogenesis*** | ***F(1,26) = 55.7*** | ***<.0001*** |
| | ***Main Effect: Training Day*** | ***F(9,234) = 12.07*** | ***<.0001*** |
| | ***Neurogenesis X Training Day*** | ***F(9,234) = 2.00*** | ***.0397*** |
